# RPE-specific MCT2 expression promotes cone survival in models of retinitis pigmentosa

**DOI:** 10.1101/2024.10.23.619878

**Authors:** Laurel C. Chandler, Apolonia Gardner, Constance L. Cepko

**Affiliations:** Departments of Genetics and Ophthalmology, Blavatnik Institute, Harvard Medical School, Boston, MA 02115; Virology Program, Harvard Medical School, Boston, MA 02115; Howard Hughes Medical Institute, Chevy Chase, MD 20815

**Author notes:** **Corresponding author:** Constance L. Cepko.

## Abstract

Retinitis pigmentosa (RP) is the most common cause of inherited retinal degeneration worldwide. It is characterized by the sequential death of rod and cone photoreceptors, the cells responsible for night and daylight vision, respectively. Although mutations in RP are mostly rod-specific, there is a secondary degeneration of cones. One possible mechanism behind cone death is metabolic dysregulation. Photoreceptors are highly metabolically active, consuming large quantities of glucose and producing substantial amounts of lactate. The retinal pigment epithelium (RPE) mediates the transport of glucose from the blood to photoreceptors and, in turn, removes lactate, which it can use as its own source of fuel. The model for metabolic dysregulation in RP suggests that, following the death of rods, lactate levels are substantially diminished causing the RPE to withhold glucose, resulting in nutrient deprivation for cones. Here, we present adeno-associated viral vector-mediated delivery of monocarboxylate transporter 2 (MCT2) into RPE cells with the aim of promoting lactate uptake from the blood and encouraging the passage of glucose to cones. We demonstrate prolonged survival and function of cones in rat and mouse RP models, revealing a possible gene agnostic therapy for preserving vision in RP. We also present the use of fluorescence lifetime imaging-based biosensors for lactate and glucose within the eye. Using this technology, we show changes to lactate and glucose levels within MCT2-expressing RPE, suggesting cone survival is impacted by RPE metabolism.

## Introduction

Retinitis pigmentosa (RP) is a group of monogenic, heritable neurodegenerative diseases that result in loss of vision for 1 in 4000 people worldwide (1). Mutant genes are primarily expressed within rods, resulting in their degeneration and the loss of night vision. As the disease progresses, there is a secondary loss of cones resulting in a deterioration of daylight color vision. Although the precise mechanisms remain unclear, various studies have suggested that inflammation (2-4), oxidative damage (5-8), and metabolic dysregulation (9-13) may all contribute to the degeneration of cones. By utilizing a gene agnostic treatment strategy to address potentially common mechanisms of cone degeneration, the therapeutic potential of such a treatment would be beneficial to a significantly larger patient population.

Rod and cone photoreceptors are highly metabolically active and rapidly metabolize glucose via aerobic glycolysis to generate substantial quantities of lactate (14-17). As photoreceptors are not vascularized, they rely on the retinal pigment epithelium (RPE) to mediate the passage of glucose from the choroid to the retina (18). Lactate is imported back across the apical membrane of the RPE through monocarboxylate transporter 1 (MCT1) and into the blood through the basolateral membrane transporter MCT3 (19). A portion of this lactate is retained by the RPE and is converted to pyruvate by lactate dehydrogenase B (LDHB), thus fueling its own metabolic needs (17). This reaction has been shown to suppress glycolysis in the RPE by depleting the available NAD^+^, thus maximizing the amount of glucose that reaches the photoreceptors (17).

The model for metabolic dysregulation in RP suggests that the loss of rods substantially reduces the level of lactate available to the RPE. As the suppressive action of lactate would be lost, the RPE may begin withholding glucose for its own glycolysis, thus depriving the surviving cones of nutrients (9, 12). We aimed to encourage the RPE to take up lactate from the blood and relinquish glucose to the surviving cones, by expressing the high affinity lactate transporter MCT2 specifically within RPE cells using an adeno-associated viral (AAV) vector. We hypothesized that the RPE would be unable to import sufficient lactate from the choroid with its basolateral transporter, MCT3, as its affinity for lactate (*K*_m_ ≈ 6mM) is too low relative to blood lactate concentrations (2.5-4.6mM) for efficient uptake from the blood (20-22). The affinity of MCT2 is, however, much higher (*K*_m_ ≈ 0.7mM) and theoretically should be able to allow the RPE to take up a greater amount of lactate within these conditions (23).

Here we demonstrate that RPE-specific AAV vector-mediated delivery of MCT2 into both rat and mouse models for RP increased cone cell survival and function across four different disease-causing mutations. We also present the use of fluorescence lifetime imaging microscopy (FLIM)-based biosensors in the eye to measure the localization, uptake, and metabolism of lactate and glucose (24, 25). We found significantly greater intracellular lactate levels and a greater accumulation of glucose in RPE cells overexpressing MCT2. Together these data suggest that increasing lactate uptake into the RPE can promote cone survival, supporting the model of insufficient or dysregulated nutrient distribution in RP as a factor in cone degeneration, and encouraging the development of therapies to address this shortcoming.

## Results

### RPE-specific MCT2 expression leads to increased cone survival in an RP rat model

To encourage glucose release and cone survival, we aimed to promote RPE lactate uptake by expressing the high affinity lactate transporter, MCT2. We generated an AAV serotype 8 vector expressing MCT2 (*Slc16a7*) under the control of the RPE-specific promoter bestrophin-1 (Best1) (AAV8.Best1.MCT2) (26). To assess the effect of RPE-specific MCT2 expression on cone survival, we initially performed paired subretinal injections in the RP rat model, S334ter line-3, which expresses a termination codon at residue S344 in the rhodopsin gene (27). Neonatal rat pups were subretinally injected with AAV8.Best1.MCT2 together with red fluorescent beads, which were used to mark the transduced region; control eyes were injected with beads only. At postnatal days 183-191 (P183-191), retinae were harvested, and surviving cones were stained using hybridization chain reaction RNA fluorescence *in situ* hybridization (HCR RNA-FISH) for retinal cone arrestin-3 (*Arr3*). By this timepoint, the rod photoreceptors will have degenerated and the outer nuclear layer reduced to less than one complete row in the most degenerated region (27); we therefore observed no residual pattern of cone survival across the retina. For this reason, we quantified cone numbers within each retina in bead positive and negative regions (Figure S1). In eyes co-injected with AAV8.Best1.MCT2, there was a statistically significant increase in the number of surviving cones in transduced relative to untransduced areas in the same eye (two-way repeated measures ANOVA, p<0.0001) (Figure 1). There was also a significant increase between the transduced regions of control and AAV8.Best1.MCT2-injected eyes (p<0.0001). There was no significant difference between the transduced and untransduced regions of control eyes (p=0.8873), nor between the untransduced regions of the control and AAV8.Best1.MCT2-injected eyes (p=0.9832). Overall, the data demonstrate that RPE-specific MCT2 expression can preserve cone survival in a rat model of RP.

**Figure 1.**
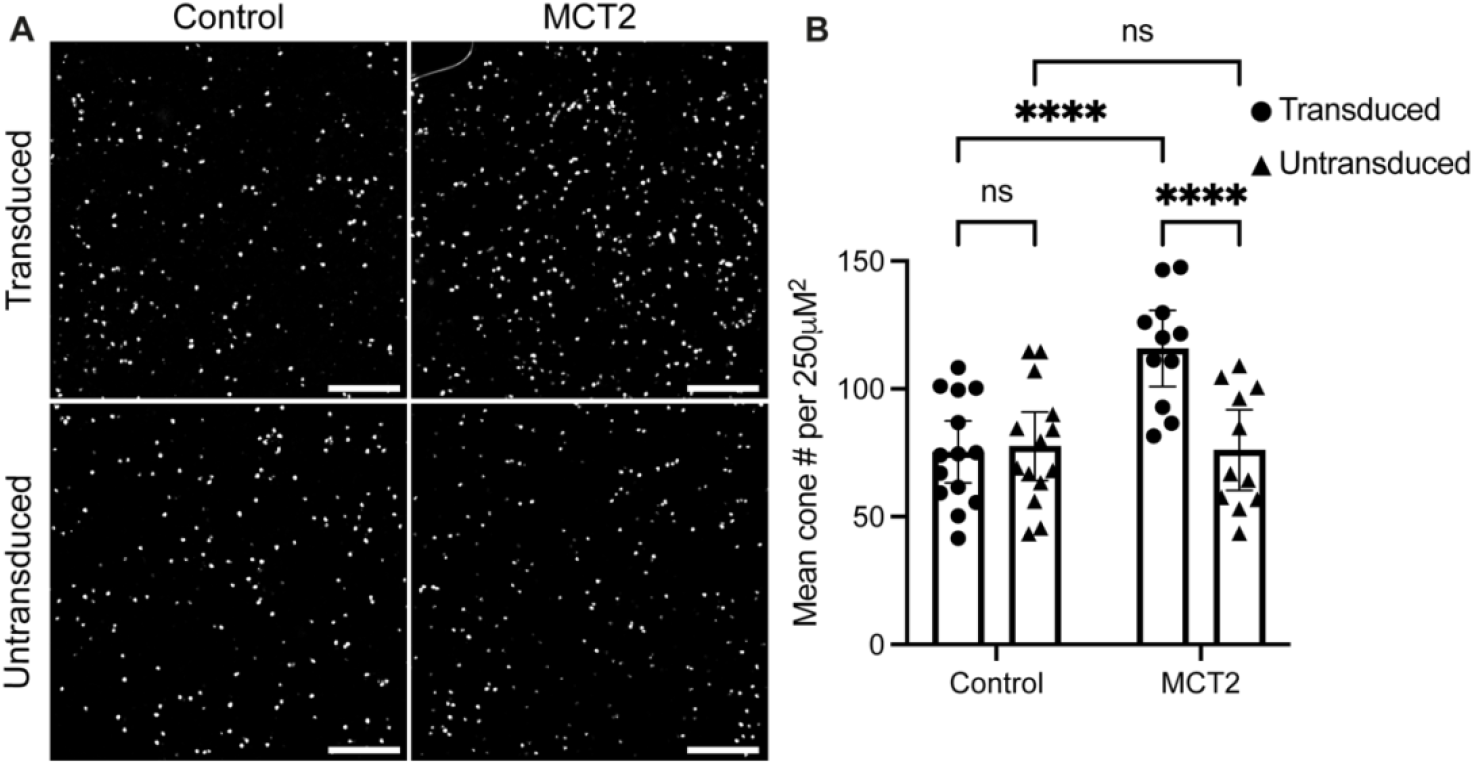
Retinal pigment epithelium (RPE)-specific monocarboxylate transporter 2 (MCT2) expression increases cone survival in a retinitis pigmentosa (RP) rat model. S334ter line-3 rats were subretinally injected with red fluorescent beads alone (control) or co-injected with AAV8.Best1.MCT2 (MCT2). Cones were identified using *in situ* hybridization against arrestin-3 (*Arr3*) at postnatal days 183-191 (P183-191). (A) Representative 500 μm^2^ images taken from the mid-periphery of transduced and untransduced regions from control and MCT2 retinae (scale bar: 100 μm). (B) Absolute cone counts averaged across four 250 μm^2^ regions within the transduced and untransduced regions of control and MCT2 samples (± 95% confidence interval (CI); control, n=14; MCT2, n=11). ****p≤0.0001 (two-way repeated measures ANOVA with Šídák’s multiple comparison test).

### RPE-specific MCT2 expression leads to increased cone survival and function in RP mouse models

We also aimed to assess the effect of RPE-specific MCT2 expression on cones in RP mouse models. Subretinal injections were performed in neonatal FVB mice, an albino mouse strain homozygous for the retinal degeneration 1 allele of *Pde6b*^*rd1*^. As the red fluorescent beads used in our rat models seemed to underestimate the transduced area, we used an AAV8 vector expressing GFP fused to histone H2B under the control of a cone-specific human red opsin (RedO) promoter to label surviving cone nuclei (AAV8.RedO.H2BGFP). Neonatal FVB mice were contralaterally injected with either AAV8.RedO.H2BGFP alone, or co-injected with AAV8.Best1.MCT2. Unlike the S334ter rat model, retinal degeneration in mice typically begins at the central retina at the optic nerve head and progresses towards the periphery; we therefore restricted our quantification to the center. When comparing GFP^+^ cone nuclei in AAV8.Best1.MCT2-injected and control retinal flatmounts at P70, there was a notable increase in the number of surviving cones in the central retina of AAV8.Best1.MCT2-injected mice (Figure 2A). Furthermore, in eyes injected with AAV8.Best1.MCT2, there was a notable absence of what appears to be “craters” or lack of cones in scattered, circular areas, a prominent feature in this FVB strain (28). Upon quantification of the number of GFP^+^ cones in the central retina, there was a significant increase in AAV8.Best1.MCT2-injected eyes compared to control (unpaired T-test, p=0.0007) (Figure 2B).

**Figure 2.**
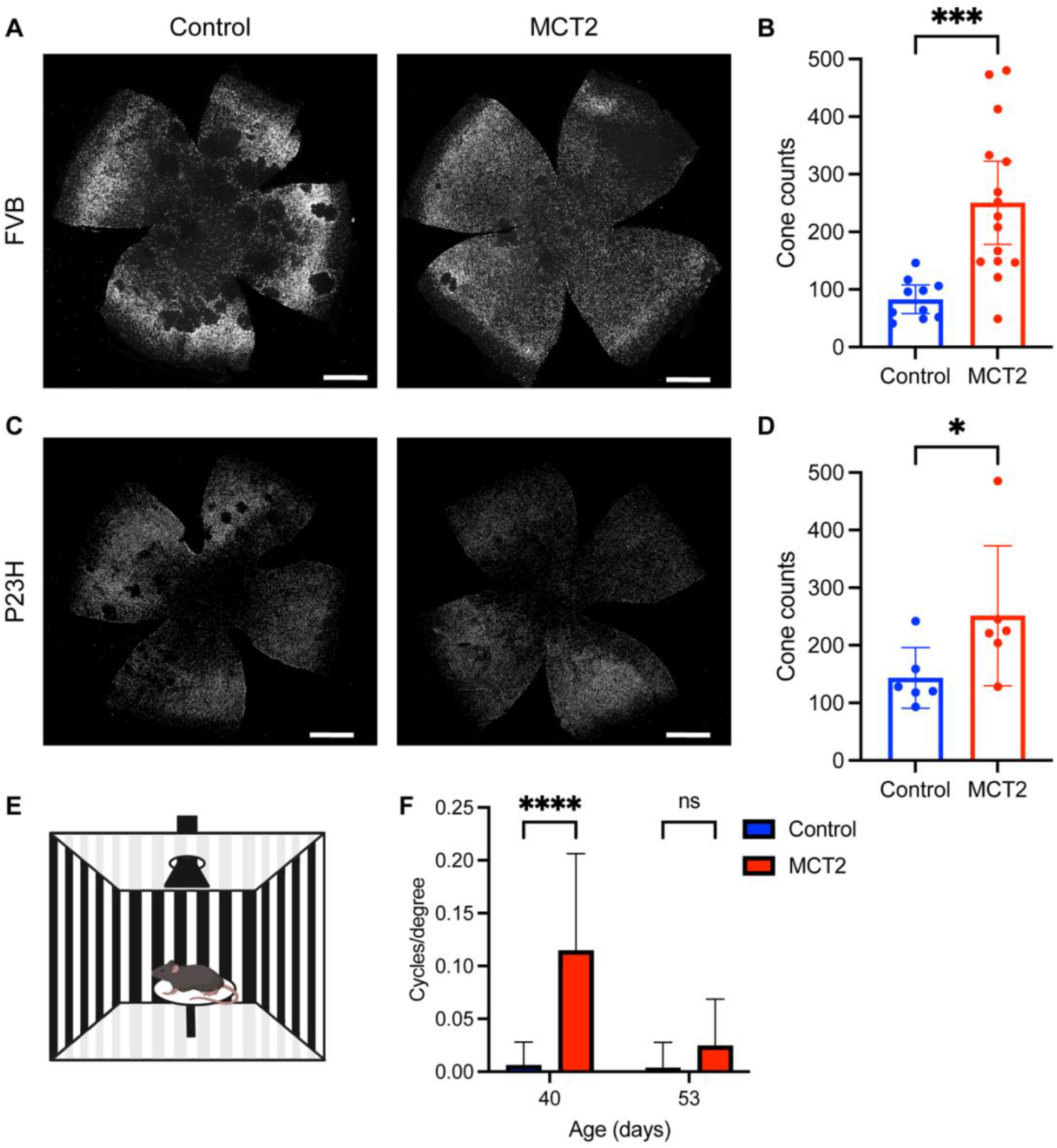
RPE-specific MCT2 expression increases cone survival and function in RP mouse models. (A&B) FVB mice were subretinally injected with AAV8.RedO.H2BGFP alone (control) or co-injected with AAV8.Best1.MCT2 (MCT2). (A) Representative image of P70 retinal flatmount with GFP^+^ cones from control or MCT2-injected eyes (scale bar: 750 μm). (B) Quantification of the number of GFP^+^ cones in the central retina (± 95% CI; control, n=10; MCT2, n=15). ***p≤0.001 (unpaired T test). (C&D) P23H mice were subretinally injected with control or MCT2. (C) Representative image of P80 retinal flatmount with GFP^+^ cones from control or MCT2-injected eyes. (D) Quantification of the number of GFP^+^ cones in the central retina (± 95% CI; n=6). *p≤0.05 (Mann-Whitney test). (E) Schematic of optomotor setup. (F) Visual acuity of paired control and MCT2 P23H eyes at P40 and P53 as measured by the number of cycles/degree using the optomotor assay (± SD; n=13). ****p≤0.0001 (two-way ANOVA with Šídák’s multiple comparison test).

The model for metabolic dysregulation suggests a common mechanism of secondary cone loss across many disease-causing mutations of RP. We therefore assessed the effect of MCT2 expression in the P23H strain, a pigmented RP mouse model with a P23H mutation in the rhodopsin gene. We subretinally injected neonatal P23H mice with either AAV8.RedO.H2BGFP alone, or co-injected with AAV8.Best1.MCT2. Retinae were harvested at P80 and demonstrated an increase in the number of central cones (Figure 2C). Furthermore, there appeared to be a small number of craters in control eyes, which were absent in AAV8.Best1.MCT2-injected retinae. Upon quantification, there was a statistically significant increase in the number of cones compared to control (Mann-Whitney test, p=0.0455) (Figure 2D). Overall, this suggests RPE-specific MCT2 expression preserves cone survival in rats and mice, with three different RP-causing mutations.

Our data demonstrate that MCT2 expression in the RPE of RP mouse models can increase survival of cone photoreceptors. However, to effectively provide vision, cone function must be preserved. We assessed cone function in RP mouse models in paired eyes injected with and without AAV8.Best1.MCT2 by measuring the optomotor reflex, which functions as a readout for visual acuity (Figure 2E). Several studies have suggested that the optomotor reflex in albino mice is absent or disrupted, therefore we were unable to assess visual acuity in FVB mice (29-31); we were also unable to assess optomotor reflex in rats due to housing constraints. We were able to assess a response in the pigmented P23H strain and, at P40, there was a significant retention of cone function in AAV8.Best1.MCT2-injected eyes compared to paired control eyes (two-way ANOVA, p<0.0001) (Figure 2F). Despite MCT2 expression preserving cone function, visual acuity continued to diminish and by P53, there was no longer a significant difference (p=0.4318). These data demonstrate that RPE-specific MCT2 expression in a RP mouse model significantly increases the duration of cone-mediated vision.

The ancillary proteins basigin and embigin have been shown to play an important role in trafficking of MCTs to the plasma membrane. Basigin has been shown to preferentially act as a chaperone for both MCT1, which is localized to the apical membrane of RPE cells, and MCT3, which is localized to the basolateral membrane (32, 33). However, the mechanism leading to their localization to opposing RPE membranes, remains unclear. The ancillary protein for MCT2 has been suggested to be embigin (34, 35). However, a recent study suggested MCT2 can be transported to the plasma membrane without a chaperone (36). To determine if MCT2 was expressed on the RPE surface membrane following AAV8.Best1.MCT2 injections, we analyzed localization in wildtype CD1 mice (Figure 3A) and degenerate FVB mice (Figure 3B) using immunohistochemistry. MCT2 expression was observed on both the apical and basal membranes of the RPE cell, demonstrating that it was transported to both cell surfaces. MCT2 was also observed in the outer plexiform layer of both control and AAV8.Best1.MCT2-injected samples, which is consistent with its expression in cone bipolar cells in a single cell RNA sequencing dataset (37).

**Figure 3.**
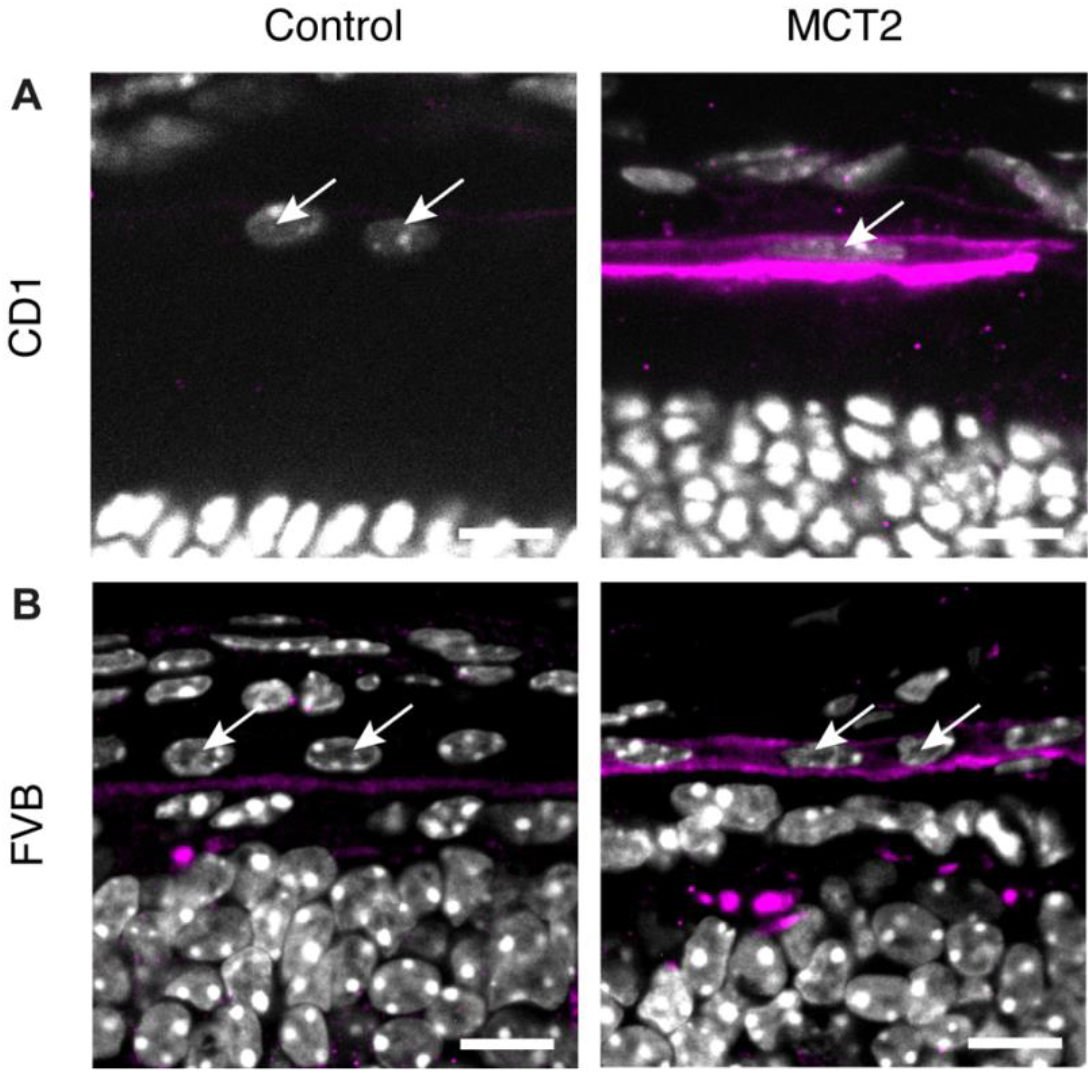
MCT2 expression on RPE cell membranes following subretinal injections with AAV8.Best1.MCT2. Representative section from (A) P42 FVB and (B) P46 CD1 mouse eyes either uninjected (control) or subretinally injected with AAV8.Best1.MCT2 (MCT2) stained with MCT2 antibody (magenta) and DAPI (white) (scale bar: 10 μm). Sections show MCT2 localization to the basolateral and apical membrane of RPE cells (RPE nuclei indicated with white arrows).

### Effect of RPE-specific MCT2 expression on the health of retinal tissue

Previously our group found that AAV vectors which express within the mouse RPE can induce toxic effects (38). We therefore aimed to determine if the AAV8.Best1.MCT2 vector induced similar changes. RPE flatmounts were stained with phalloidin, which labels F-actin in the cell membrane and reveals the hexagonal shape of healthy RPE cells. In wildtype Sprague Dawley rats only minor changes to RPE morphology were observed (Figure S2), suggesting that RPE-specific expression of MCT2 did not induce significant toxic effects.

Although the AAV8.Best1.MCT2 vector did not induce significant toxicity in rats, there did appear to be some toxic effects in mice. At P40, FVB mouse eyes subretinally injected with AAV8.Best1.MCT2 were harvested for staining of the RPE with phalloidin. In control eyes, the RPE showed signs of disruption due to progressing retinal degeneration (Figure S3). However, in AAV8.Best1.MCT2-injected eyes, there were additional morphological changes consistent with RPE toxicity, including the enlargement of cells and the presence of stress fibers (Figure S3) (38). This observation indicates that mice were more sensitive to RPE-specific AAV expression than rats, despite receiving a lower dose than rats. Nonetheless, there were significant improvements in cone survival and function in mice.

### Metabolic changes in the MCT2-expressing RPE

The rationale behind delivering MCT2 into RPE cells was to increase lactate uptake and prevent retention of glucose. To probe this mechanism, we utilized FLIM-based biosensors that specifically detect lactate and glucose. Fluorescence lifetime is the average time between photon excitation and emission, and, when used together with two-photon technologies, can robustly quantify the levels of a given molecule within a tissue. The benefit of lifetime measurements compared to fluorescence intensity readouts is that these measurements are independent of biosensor expression level and have reduced pH sensitivity (39). We generated AAV vectors expressing the FLIM-based lactate sensor LiLac and the glucose sensor GlucoSnFR-TS under the control of the RPE-specific Best1 promoter (24, 25). The LiLac sensor is composed of the lactate binding protein TlpC from *Helicobacter pylori*, which is inserted within the mTurquoise2 fluorophore sequence with the N- and C-termini of TlpC connected by flexible linkers (24). TlpC can generate a large conformational change upon binding to lactate which in turn causes lifetime shifts in the mTurquoise2 fluorescent protein. The bound version of LiLac has a shorter lifetime value compared to the unbound. In contrast, GlucoSnFR-TS is a T-sapphire fluorescent protein inserted between residues 326 and 327 of *Thermus thermophilus* glucose binding protein and has a longer lifetime value when bound (25).

Initially, to assess the levels of lactate within MCT2-expressing RPE, we performed subretinal injections of AAV8.Best1.LiLac with and without AAV8.Best1.MCT2 into neonatal FVB mice. Three weeks later, the eyecup was harvested, flattened, and immediately imaged using a two-photon fluorescent lifetime microscope to visualize lifetime changes in the RPE. The RPE tissue was incubated in Kreb’s ringer bicarbonate (KRB) buffer with increasing concentrations of lactate (0, 0.1, 1, and 5 mM) that reflect the approximate blood lactate concentrations in mice (2.5-4.6 mM) (21, 22). At baseline, AAV8.Best1.MCT2-injected RPE samples had a significantly shorter lifetime compared to control, demonstrating that MCT2-expressing RPE cells have greater intracellular lactate levels (two-way ANOVA, p<0.0001) (Figure 4A and B). When incubated with increasing concentrations of lactate, both control and AAV8.Best1.MCT2-injected samples had no detectable change at 0.1 mM compared to baseline (p=0.7257 (control), p=0.2901 (MCT2)). However, both showed shorter lifetimes at 1 and 5 mM lactate, suggesting lactate uptake into these cells (p<0.0001 for all conditions). AAV8.Best1.MCT2-injected samples also had significantly shorter lifetimes at each lactate concentration compared to control (p<0.0001 for all conditions), suggesting that, despite having greater lactate levels at baseline, RPE cells expressing MCT2 continued to import lactate from the media.

**Figure 4.**
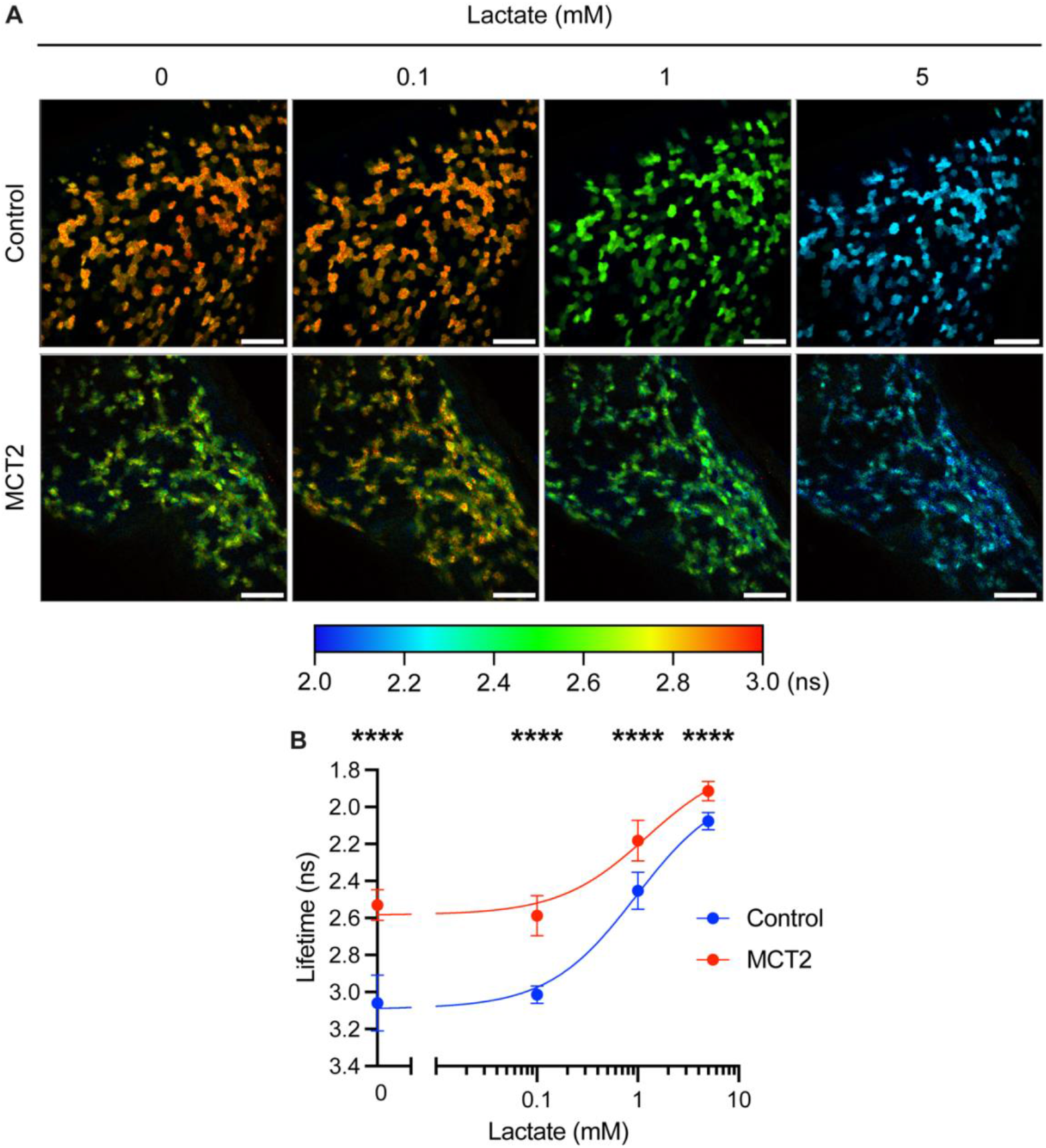
MCT2 overexpression leads to increased intracellular lactate in the RPE. Neonatal FVB mice were subretinally injected with AAV8.Best1.LiLac alone (control) or co-injected with AAV8.Best1.MCT2 (MCT2). At P20-21, eyecups (RPE, choroid, and sclera) were harvested and RPE cells imaged *en face* using two-photon fluorescent lifetime imaging microscopy (FLIM). (A) Representative lifetime images from control and MCT2 samples sequentially incubated with 0, 0.1, 1, or 5 mM lactate. Color scale represents increasing lifetime in nanoseconds (ns), which corresponds to decreased intracellular lactate concentrations (scale bar: 100 μm). (B) Lifetime at each lactate concentration (± SD; control, N_mice_=4; N_regions_=12; MCT2, N_mice_=6, N_regions_=16). Y-axis reversed to reflect rise in lactate concentration with decreasing lifetime and data fit with a dose-response curve. ****p≤0.0001 (two-way ANOVA with Šídák’s multiple comparison test).

Next, we aimed to assess glucose levels as a readout for glucose uptake and/or glycolysis. Neonatal FVB mice were subretinally injected with AAV8.Best1.GlucoSnFR-TS with and without AAV8.Best1.Mct2. Three weeks later, the RPE tissues were processed and incubated with 0, 2, 5, and 10 mM glucose in KRB buffer, which reflects the approximate blood glucose levels of FVB mice (9.1 mM) (40). At baseline, with 0 mM glucose, there was no significant difference between the lifetime in control and AAV8.Best1.MCT2-injected samples, indicating that intracellular glucose levels were comparable (two-way ANOVA, p=0.9998) (Figure 5A and B). However, upon incubation in increasing concentrations of glucose, the lifetime was significantly longer in the AAV8.Best1.MCT2-injected samples, showing greater intracellular concentrations of glucose (p<0.0001 (2 and 5 mM), p=0.0138 (10 mM)) (Figure 5A and B). As differences in intracellular glucose concentration could be due to differences in uptake and/or the rate of glycolysis, we sought to determine if a reduction in glycolysis in this tissue under these culture conditions would lead to greater intracellular glucose. To test this, we pre-incubated the RPE samples for 10 minutes with 0.5 mM of the GAPDH inhibitor iodoacetic acid (IAA) to block glycolysis. At 0 mM glucose there was no statistically significant difference in either the control or AAV8.Best1.MCT2-injected samples following IAA incubation, suggesting that, without added glucose, there was little glycolysis being performed (one-way ANOVA, p=0.4642) (Figure 6A). Following incubation with 10 mM glucose, there was a statistically significant increase in the fluorescent lifetime in both control (one-way ANOVA, p<0.0001) and AAV8.Best1.MCT2-injected samples (p=0.0003) (Figure 6B and C). This suggests that the *ex vivo* RPE tissue cultures used in this study can import and metabolize glucose, allowing for the hypothesis that an increased intracellular concentration of glucose can be due to decreased glycolysis.

**Figure 5.**
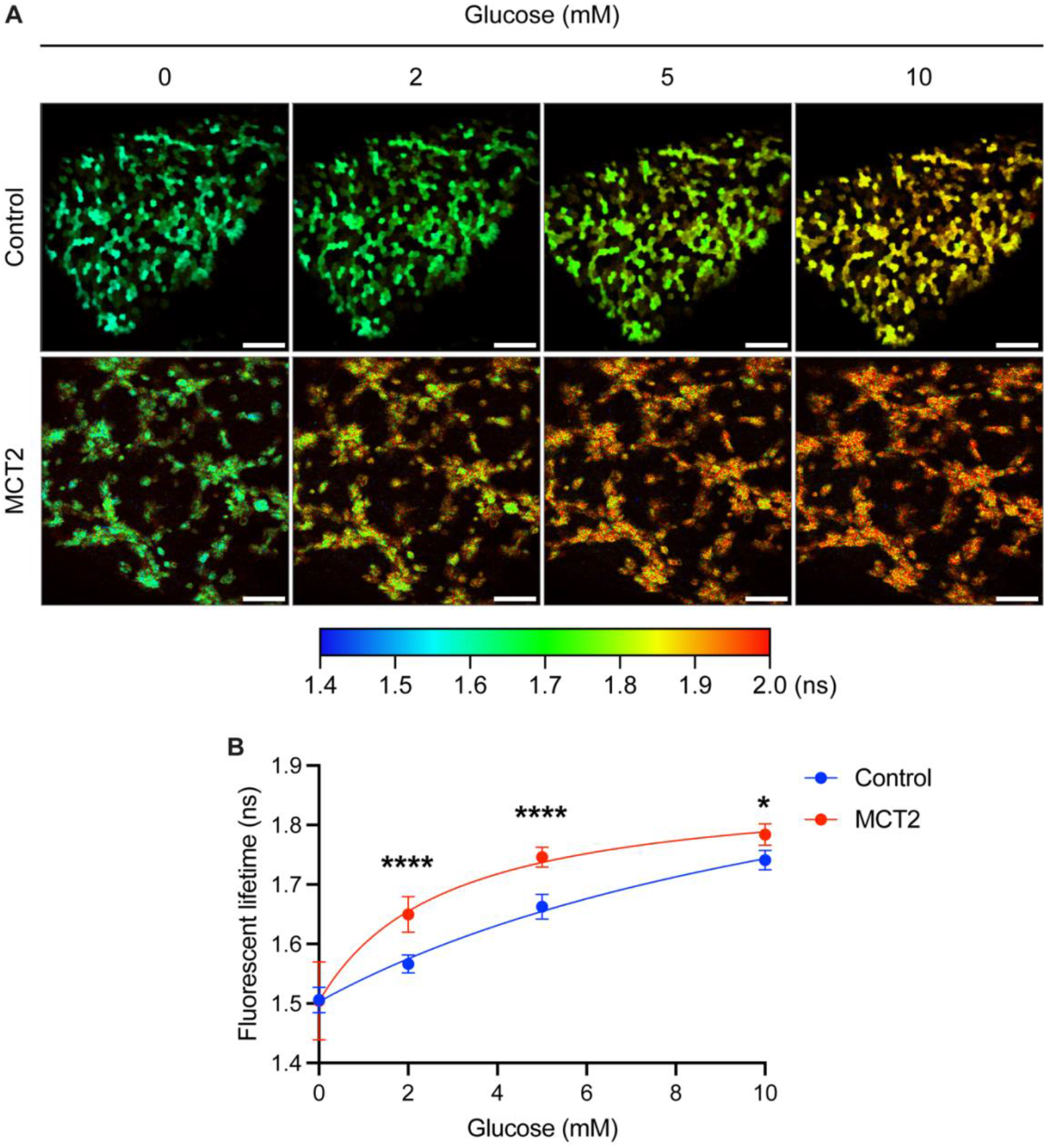
MCT2 overexpression leads to increased intracellular glucose in the RPE. Neonatal FVB mice were subretinally injected with AAV8.Best1.GlucoSnFR-TS alone (control) or co-injected with AAV8.Best1.MCT2 (MCT2). At P20-21, eyecups were harvested and RPE cells imaged *en face* using two-photon FLIM. (A). Representative lifetime images from control and MCT2 samples sequentially incubated with 0, 2, 5, and 10 mM glucose. Color scale represents increasing lifetime in ns, which corresponds to increased intracellular glucose concentrations (scale bar: 100 μm). (B) Lifetime at 0, 2, 5, and 10 mM glucose (± SD; N_mice_=3; N_region_=9). Data fit with a dose-response curve. *p≤0.05, ****p≤0.0001 (two-way ANOVA with Šídák’s multiple comparison test).

**Figure 6.**
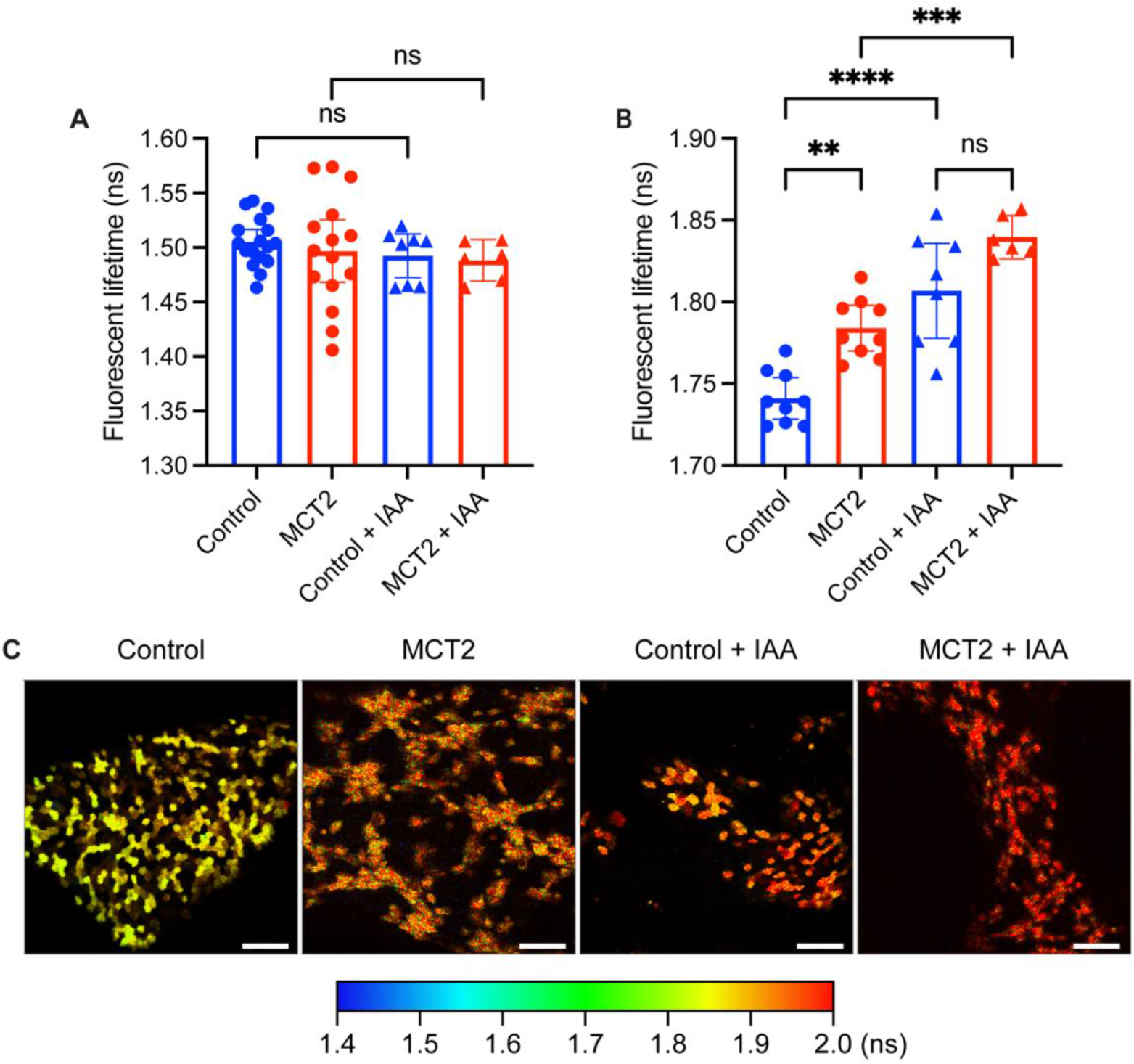
Inhibition of glycolysis increases intracellular glucose concentrations in the RPE. Neonatal FVB mice were subretinally injected with AAV8.Best1.GlucoSnFR-TS alone (control) or co-injected with AAV8.Best1.MCT2 (MCT2). At P20-21, eyecups were harvested and RPE cells imaged *en face* using two-photon FLIM. Following baseline lifetime recordings, the same RPE samples were treated with 0.5 mM iodoacetic acid (IAA) for 10 minutes. (A) Lifetime at baseline (0 mM glucose) with and without IAA treatment (± 95% CI; control, n=17; MCT2, n=15; control + IAA, n=8; MCT2 + IAA, n=6). (B) Lifetime of RPE incubated with 10 mM glucose following treatment with or without IAA (± 95% CI; control, n=9; MCT2, n=9; control + IAA, n=8; MCT2 + IAA, n=6). (C) Representative images from control and MCT2 samples incubated with 10 mM glucose and treated with and without IAA (scale bar: 100 μm). **p≤0.01, ***p≤0.001, ****p≤0.0001 (one-way ANOVA with Tukey’s multiple comparison test).

## Discussion

The current model for metabolic dysregulation in RP suggests that retinal lactate levels are substantially diminished due to the death of rods. This in turn causes the RPE to sequester glucose for its own metabolic needs and ultimately results in the starvation and degeneration of cones (9, 12). In this study we delivered the high affinity lactate transporter MCT2 to the RPE with the aim of promoting lactate uptake from the underlying choroidal vasculature and encouraging the release of glucose. Subretinal injections of an AAV8.Best1.MCT2 vector in rat and mouse models of RP promoted cone survival across several disease-causing mutations, as well as cone function. However, despite MCT2-expression increasing retention of cone function, this effect diminished with time, something which has been noted with other cone-rescuing therapies (3, 7, 13). It is likely that other mechanisms also contribute towards cone degeneration, including inflammation (2-4) and oxidative damage (5-8). To achieve sustained rescue, each of these may need to be addressed. Nevertheless, these data support the hypothesis that the metabolic status of the RPE impacts cone survival and reveals a novel gene-agnostic therapy for prolonging vision in RP. Furthermore, a genome-wide association study found a splice variant of MCT3 (*Slc16a8*) was correlated with age-related macular degeneration (41), while another found AAV-mediated overexpression of MCT2 in retinal ganglion cells of a glaucoma mouse model resulted in increased cell survival and metabolic improvements (42). Together this demonstrates disrupted monocarboxylate transport is associated with retinal diseases and presents a possible therapeutic benefit for increased uptake.

A lack of appropriate techniques to assess the localization and levels of metabolites within the retina has made it challenging to investigate the mechanism of therapies that target metabolic dysregulation. The current techniques available to measure intracellular metabolite levels have been mainly limited to whole tissue analysis with a lack of cell-type specificity or single cell resolution. In this study, we developed the use of FLIM-based biosensors within the eye, which enables cell-specific measurement of lactate and glucose localization and concentration. Using the FLIM-based biosensor for lactate, LiLac, we detected significantly greater intracellular lactate levels in AAV8.Best1.MCT2-injected RPE cells. This finding supports our proposed mechanism for AAV8.Best1.MCT2-mediated cone survival. It is possible that reduced lactate metabolism in these cells could also lead to increases in intracellular concentrations of lactate. However, due to the function of MCT2 and its detection on the cell surface, it is likely that an increase in uptake resulted in these differences.

With the use of the FLIM-based biosensor for glucose, GlucoSnFR-TS, we did not detect a difference in intracellular glucose levels at baseline. However, incubation with supplementary glucose resulted in a greater amount of glucose within MCT2-expressing RPE cells, suggesting reduced glycolysis and/or increased uptake. Since MCT2 expression was unlikely to directly affect glucose uptake, we hypothesized that such an effect was due to a reduction in the rate of glycolysis. Treatment with the glycolysis inhibitor IAA, in conjunction with 10 mM glucose, resulted in a significant accumulation of intracellular glucose in both control and AAV8.Best1.MCT2-injected samples. The same MCT2-expressing RPE cells were seen to have increased intracellular lactate, which has been shown to inhibit glycolysis (17). Together, these findings are consistent with the hypothesis that MCT2-expressing RPE cells have higher intracellular glucose due to reduced rates of glycolysis, but do not rule out that they may have a different rate of glucose uptake, instead of, or in addition to, reduced glycolysis. Since the first step of glycolysis results in the irreversible retention of glucose, these findings support the model that cone degeneration is due to the RPE switching to glycolysis and withholding glucose. IAA treatment, however, had no detectable effect on lifetimes in control or AAV8.Best1.MCT2-injected samples at baseline, suggesting that neither sample was performing glycolysis. Since RPE samples could perform glycolysis when glucose was present, this suggests a lack of intracellular glucose. GlucoSnFR-TS is highly specific to glucose and will not bind to any glycolytic intermediates, therefore the lack of detectable glucose at baseline may suggest residual levels are rapidly metabolized following tissue harvest (43). For this reason, the use of *ex vivo* tissue has its limitations. Although we analyzed tissue as quickly as possible after harvesting to prevent substantial changes, the process of taking the tissue out of the whole organism will likely impact metabolism and may lead to depletion of some metabolites. Live imaging of the eye with adaptive optics fluorescent lifetime imaging ophthalmoscopy (AOFLIO) in conjunction with these FLIM-based biosensors, could enable a more accurate assessment of retinal metabolism *in vivo* (44, 45). Nonetheless, this technique demonstrates a significant improvement in the ability to detect metabolites in the retina. Overall, these data support our hypothesis for increased lactate uptake and reduced glycolysis in RPE cells as a possible therapy to support cone survival in RP.

Our aim in delivering MCT2 to RPE cells was for it to be expressed on the basolateral membrane to promote lactate uptake from the underlying choroidal vasculature. Although we were able to detect MCT2 on this membrane, we also detected it on the apical membrane. The lower affinity transporter MCT1, which is also expressed on the apical membrane, functions to take up lactate produced by photoreceptors (19). The implication of MCT2 expression on the apical membrane is unclear and it may encourage additional uptake of lactate from the subretinal space. Furthermore, in addition to lactate, members of the MCT family can import other monocarboxylates including pyruvate and ketone bodies (33). Ketone bodies are produced by the RPE following metabolism of fatty acids acquired from phagocytosis of photoreceptor outer segments (46). These ketone bodies are exported from the cell through MCT1 and can be taken up by photoreceptors and oxidized to carbon dioxide (19, 46). As MCT2 can also transport ketone bodies, it is possible that this has an additional effect on the metabolism of RPE and cones. Furthermore, the RPE does not solely rely on lactate as its source of nutrients, with several studies suggesting other potential fuel sources, including fatty acids (46, 47), glutamine (48), proline (49), and succinate (50, 51). Although the data presented in this study suggests a possible role for lactate in RPE metabolism, further investigation into the other metabolites during cone degeneration should be explored.

We previously published that RPE-expressing AAV vectors induce ocular toxicity in mice (38). Interestingly, although we detected toxicity in mouse models subretinally injected with AAV8.Best1.MCT2, we did not observe significant toxicity in rats. As the rat eye is significantly larger than the mouse, it is possible that mice received a larger number of viral genomes per cell and, due to the dose-dependent nature of AAV-induced toxicity, larger doses would result in greater levels of toxicity (38). Nonetheless, we saw changes in RPE morphology in mice at doses as low as 2.5×10^7^ genome copies (gc)/eye (data not shown), while minimal changes were seen at a dose of 2×10^9^ gc/eye in rats. Since the rat eye is only approximately double the size of a mouse eye, this 80-fold difference in dose is unlikely to account for the observed differences (52). These changes may be explained by a mouse-specific sensitivity to the overexpression of transgenes within the RPE. Although the mechanism behind this RPE toxicity remains unclear, there are several immune mechanisms lacking in mouse which are seen in both rats and humans, such as a lack of major histocompatibility complex (MHC) class II molecules on T cells (53). Furthermore, we cannot rule out that MCT2 itself could be contributing to these morphological changes. MCT1-4 all couple lactate transport with a proton (33, 54), therefore changes in the expression of these transporters may alter intracellular pH and induce toxicity. For instance, in one study the knockout of MCT3 in mice was associated with reduced electroretinogram (ERG), and in another, the RPE-specific knockout of basigin, which functions as a chaperone for MCT1 and MCT3 in their transport to the cell surface, induced reductions in ERG responses, disruption to photoreceptors, and changes to RPE morphology (55, 56). It is therefore possible that overexpression of MCT2 may be responsible for the mouse-specific toxicity. A means of assessing its effect on intracellular pH could be achieved using the genetically encoded pH sensor, pHRed, to detect lifetime changes and measure relative pH differences within MCT2-expressing RPE cells (57). Together, this raises the question about the choice of animal model when assessing both the efficacy and safety of potential therapeutics.

Overall, we demonstrate that AAV vector-mediated expression of MCT2 in RPE cells of rat and mouse models of RP improves cone survival and function across a range of mutations. We also present the use of FLIM-based biosensors in the eye that help uncover the mechanism of MCT2-mediated cone survival. The data suggest that this transporter is promoting lactate uptake and reducing retention of glucose. The development of FLIM-based biosensors for use within the eye may help elucidate the mechanism of other treatments targeting metabolic imbalances in RP and ultimately inform the design of future therapeutics. Together this supports the model for changes to RPE metabolism having an impact on cone survival in RP.

## Materials and Methods

### Animals

S334ter line-3 rats (strain #00643) were purchased from Rat Resource & Research Center, University of Missouri (27). Sprague Dawley rats (strain #001) were purchased from Charles River Laboratories. FVB (strain #207) and CD1 mice (strain #022) were purchased from Charles River Laboratories. P23H mice (strain #017628) were purchased from The Jackson Laboratory. Animals were bred and maintained at Harvard Medical School on a 12-hour alternating light and dark cycle. All experimental procedures were approved by the Institutional Animal Care and Use Committee at Harvard University. Both males and females were used in all experiments and randomly assigned to groups.

### Vector design

The vector AAV8.Best1.MCT2 was generated by replacing the GFP sequence in the AAV.Best1.GFP plasmid, which had been created by a fellow lab member, Wenjun Xiong (26, 38), with the mouse *Slc16a7* (MCT2) cDNA sequence. RNA was isolated from mouse retinae using the miRNeasy Mini Kit (QIAGEN) according to manufacturer’s instructions. The RNA was reverse transcribed into cDNA using SuperScript® III First-Strand Synthesis Kit (Invitrogen) and purified using the QIAquick PCR Purification Kit (QIAGEN) according to manufacturer’s instructions. The *Slc16a7* sequence was amplified using PCR and inserted into the AAV.Best1.GFP plasmid using Gibson cloning. AAV8.RedO.H2BGFP expressed a GFP fused to H2B and was under the control of a cone-specific RedO promoter (13, 28, 58, 59); this promoter was a gift of the Botond Roska laboratory, Friedrich Miescher Institute for Biomedical Research, Basel, Switzerland (60). The vectors AAV8.Best1.LiLac and AAV8.Best1.GlucoSnFR-TS were generated from plasmids gifted by the Gary Yellen laboratory (24, 25), Harvard Medical School and cloned into Best1 promoter plasmids (26). All AAV vectors were single stranded.

### AAV Production

AAV vectors were produced as previously described (7, 61). HEK293T cells were transfected with a total of 170 μg of plasmid DNA using polyethylenimine in DMEM supplemented with 10% NuSerum and 1% penicillin/streptomycin. The vector plasmids were co-transfected with the Rep2/Cap8 packaging plasmid (pAAV2/8) and the adenoviral helper plasmid (pHGTI.adeno1). 24 hours post-transfection, the media was changed to serum-free DMEM supplemented with 1% penicillin/streptomycin. 72 hours post-transfection, the media was harvested and precipitated, and the cells were harvested and lysed. The AAV particles were isolated by ultracentrifugation with an iodixanol gradient, purified using Amicon Ultra-15 100K filter units (MilliporeSigma), and collected in sterile PBS with 0.0001% Pluronic F-68 (Thermo Fisher Scientific). The viral titers were semi-quantitatively determined by staining an SDS-PAGE with SYPRO Ruby Protein Gel Stain (Invitrogen) to visualize the VP1, VP2, and VP3 viral capsids relative to a reference vector with a known titer.

### Subretinal injections

Subretinal injections were performed in neonatal mice at P0-2. Pups were anesthetized on ice and were administered 0.05-0.1 mg/kg buprenorphine immediately prior to the subretinal injection, as well as a drop of proparacaine hydrochloride ophthalmic solution in each eye post-subretinal injection. The palpebral fissure was cut with a 30 G needle, the eyelids were opened, and the eye was gently proptosed with blunt forceps. Injections were performed using glass needles made by pulling Wiretrol II capillaries on a needle puller (Sutter Instruments, Model P-97) and beveled with a microgrinder (Narishige). The needles were connected to a FemtoJet Microinjector (Eppendorf) and gently inserted into the subretinal space through the sclera, choroid, and RPE for injection. In rats, 2×10^12^ gc/mL AAV8.Best1.MCT2 was co-injected with 0.01% red fluorescent beads (0.1 μm red FluoSpheres™, excitation/emission 580/605, Invitrogen) and 0.1% FastGreen (Millipore Sigma), in approximately a 1 μL volume per eye. For all rats, right eyes were injected with beads only and left eyes were co-injected with AAV8.Best1.MCT2. In mice, AAV vectors were co-injected with 0.1% FastGreen, in approximately a 0.25 μL volume per eye. For all histology experiments performed in mice, 4×10^11^ gc/mL of AAV8.Best.MCT2 was delivered. For the P23H optomotor experiments 1×10^11^ gc/mL of AAV8.Best1.MCT2 was delivered. For FLIM experiments, 4×10^11^ gc/mL of AAV8.Best.MCT2 and 1×10^12^ gc/mL of either AAV8.Best1.LiLac or AAV8.Best1.GlucoSnFR-TS was delivered. In experiments using AAV8.RedO.H2BGFP 1×10^12^ gc/mL of vector was delivered.

### Histology

Eyes to be used for cone counting were enucleated and dissected in PBS to remove the retina. For mouse retinae, tissue was fixed in 4% paraformaldehyde (PFA) for 30 minutes at room temperature and then washed three times in PBS prior to mounting with Fluoromount-G mounting medium (Invitrogen). For rat retinae, the tissue was used for HCR RNA-FISH.

For retinal sections, eyes were dissected to remove the cornea, iris, and lens and leave the eye cup intact. Eyecups were cryoprotected by incubating in 7.5% sucrose until equilibrated before transferring to a solution of 50% OCT compound and 15% sucrose for at least 30 minutes. Eyecups were flash frozen in this OCT/sucrose mixture and stored at -80°C prior to sectioning with a Leica CM3050S cryostat (Leica Microsystems). Sections were hydrated by washing three times with PBS before incubating in LAB buffer (PolySciences Ltd) for 5 minutes at room temperature for a gentle antigen retrieval. Sections were washed once in PBS before incubating with blocking buffer (4% donkey serum, 0.3% triton x-100, 0.3% bovine serum albumin (BSA)) for one hour at room temperature. Sections were incubated with a primary antibody against MCT2 (Proteintech; 20355-1-AP) at a 1/200 dilution in blocking buffer overnight at 4°C in a humidified chamber. The following day, sections were washed three times with PBS. The tissue was incubated with anti-rabbit 647 secondary antibody (Jackson ImmunoResearch) at a 1:750 dilution in blocking buffer for two hours at room temperature in the dark. Sections were washed three times in PBS and stained with DAPI (Invitrogen) for five minutes before three subsequent washes. Sections were mounted in Fluoromount-G mounting medium.

For RPE flatmounts, after enucleation and removal of the cornea, iris and lens, the retina was carefully removed to expose the apical surface of the RPE, attached to the choroid and sclera. The remaining RPE eyecup was fixed in 4% PFA for two hours at room temperature, followed by three washes with PBS. The eyecups were incubated with 1/100 phalloidin Alexa fluor 647 (Thermo Fisher Scientific) in blocking buffer (1% Triton-X and 1% BSA in PBS) overnight at 4°C. The eyecups were washed three times in PBS and flattened with radial cuts and mounted onto a cover slip and slide.

### HCR RNA-FISH

Rat retinae were stained using a protocol adapted from manufacturer’s instructions with probes to *Arr3* (Molecular Instruments) (62). After fixation and washing in PBS, retinae were permeabilized in 70% ethanol for two hours at room temperature. Samples were washed in probe wash buffer preheated to 37°C for five minutes at room temperature, and then incubated in hybridization buffer at 37°C for 30 minutes. A probe solution was prepared by incubating *Arr3* probes (1 μM stock) in hybridization buffer to a final concentration of 60 nM and was pre-heated to 37°C for 30 minutes. The retinae were subsequently incubated in the probe solution at 37°C overnight. The next day, the retinae were washed twice in probe wash buffer at room temperature for 30 minutes each. This was followed by two washes at room temperature in 100% 5X SSC buffer with 0.1% Tween (SSCT buffer) for 20 minutes each. The H1 and H2 hairpin B1 647 nm amplifiers (3 μM stock) were heated to 95°C for 90 seconds and then cooled to room temperature for 30 minutes in the dark. The hairpins were combined with probe amplification buffer at a final concentration of 48 nM. The retinae were pre-incubated with amplification buffer for 10 minutes before incubating overnight in the hairpin mixture in the dark at room temperature. The following day, the samples were washed twice in 5X SSCT buffer for 20 minutes at room temperature before flattening onto a coverslip and mounting.

### Image acquisition and analysis

All retinal flatmount images and the rat phalloidin stained RPE were acquired on an Olympus VS200 Slide Scanner with a UPlan X Apo 10x/0.4 Air objective. FVB phalloidin stained RPE was acquired on a Nikon Ti inverted microscope with a W1 Yokogawa Spinning disk with 50 um pinhole disk and a Plan Apo λ 20x/0.75 DIC I objective. FVB and CD1 sections were imaged on the Nikon Ti2 inverted microscope with a W1 Yokogawa Spinning disk with 50 um pinhole disk and a Plan Apo λ 60x/1.4 Oil DIC objective.

GFP^+^ cone nuclei infected with AAV8.RedO.H2BGFP were imaged using the 488 laser and quantified using a modified version of an ImageJ macro as previously described (3, 4). In rats, we defined the transduced and untransduced area based on signal from the red fluorescent beads and the macro assigned four 250 μm^2^ polygons at the midpoint between the optic nerve and the outer retinal boundary in each of these regions (Figure S1). The cone count was the mean of these four regions. For mice, the same macro was used as previously described (3, 4), however, to limit the quantification to only the central retina, the region where the cones were counted was generated as ¼ of the distance from the optic nerve to the periphery rather than ½.

### Optomotor assay

Optomotor assays were performed using the OptoDrum (Straitech, Germany) automated system to determine the number of degrees per cycle. Mice were contralaterally injected with either AAV8.Best1.MCT2 and AAV8.RedO.H2BGFP, or AAV8.RedO.H2BGFP alone. All animals were habituated on the center platform for five minutes prior to the start of the first experiment. The black and white stripes were displayed at 99.7% contrast with the rotation speed of 12 degrees/second in both a clockwise or counterclockwise direction to assess the left and right eye, respectively. The OptoDrum software determined whether a mouse performed the corresponding head movement. The frequency of the stripes (cycles/degree) gradually increased until the mice failed to track three times.

### Two-photon FLIM

Mice aged P20-21 were euthanized, the eye was enucleated in ice cold HBSS and the cornea, iris, and lens removed. One radial incision was made in the remaining eyecup and the retina was carefully removed to expose the RPE. An additional seven radial incisions were made, and the eyecup was transferred to KRB buffer (98.5 mM NaCl, 4.9 mM KCl, 2.6 mM CaCl2, 1.2 mM MgSO4, 1.2 mM KH2PO4, 26 mM NaHCO3, 20 mM HEPES) before immediately proceeding with imaging (63, 64). The eyecup was transferred to a coverslip with the apical membrane of the RPE facing up. Residual buffer was wicked away using a Kimwipe to enable the eyecup to be gently flattened and the tissue was kept flat by using a slice anchor (Warner Instruments). The coverslip was transferred to the custom-made perfusion chamber pre-filled with KRB buffer at room temperature. Lactate or glucose was delivered in KRB buffer through the perfusion system and samples were incubated for five minutes before data acquisition. Following IAA delivery, samples were incubated for 10 minutes.

Lifetime imaging was performed with a Leica upright DM6 Stellaris 8 Dive Falcon and a Plan Apo 25x/0.9 immersion objective. For LiLac, the light source was a two-photon tunable Insight X3 dual beam laser tuned to 850 nm at 8% laser intensity with a gain of 10 and emission light was bandpass filtered (465-550 nm). For GlucoSnFR-TS, the laser was tuned to 790 nm at 5% laser intensity with a gain of 10. For LiLac, 12 line accumulations were acquired for control samples and 15 for AAV8.Best.MCT2-injected samples. For GlucoSnFR-TS, 10 line accumulations were acquired for control and 10-15 for AAV8.Best.MCT2-injected samples. Phasor analysis was performed using the LAS X software (Leica) on data acquired from binning multiple Z stacks. The wavelet filter was applied to the phasor and the lifetime was measured at the center of the population (Figure S4).

A dose-response curve was fitted with a Hill equation (equation 1) where data was plotted as lifetime (LT) against concentration of lactate or glucose (conc) (24, 25). LT_min_ and LT_max_ correspond to the lower and upper plateau, respectively, K_0.5_ is the concentration at the halfway point between LT_min_ and LT_max_, and n_H_ is the Hill coefficient, which was set to -1.0 for LiLac and 1.0 for GlucoSnFR-TS.

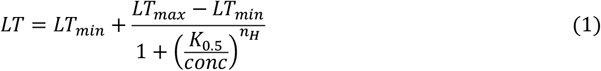

### Statistics

All statistical analysis was performed using GraphPad Prism Version 10. Datasets were tested for normality using a Shapiro-Wilk test. Data that was normally distributed was analyzed using a parametric test (T-test or ANOVA), while skewed data were analyzed using a non-parametric test (Mann-Whitney test). Multiple comparisons were corrected for using a Šídák’s or Tukey’s multiple comparison test for two- or one-way ANOVA, respectively. All statistical analyses were performed using an alpha of 0.05.

## Acknowledgments

We would like to thank Professor Gary Yellen, Paul Rosen, and Professor Carlos Manlio Diaz Garcia for their invaluable advice with the FLIM biosensors and for kindly providing us with the plasmids for LiLac and GlucoSnFR-TS. We would also like to thank Dr Sophia Zhao for generating the AAV8.RedO.H2BGFP virus and for Lucas Lin for his help performing optomotor assays. We would finally like to thank the MicRoN microscopy core at Harvard Medical School for their expert microscopy advice. This study was funded by Howard Hughes Medical Institute (C.L.C), Spark Therapeutics (L.C.C and A.G), the Harvard Brain Institute (LCC), and NIH (A.G).

