## Supplemental Figures for "RPE-specific MCT2 expression promotes cone survival in models of retinitis pigmentosa"

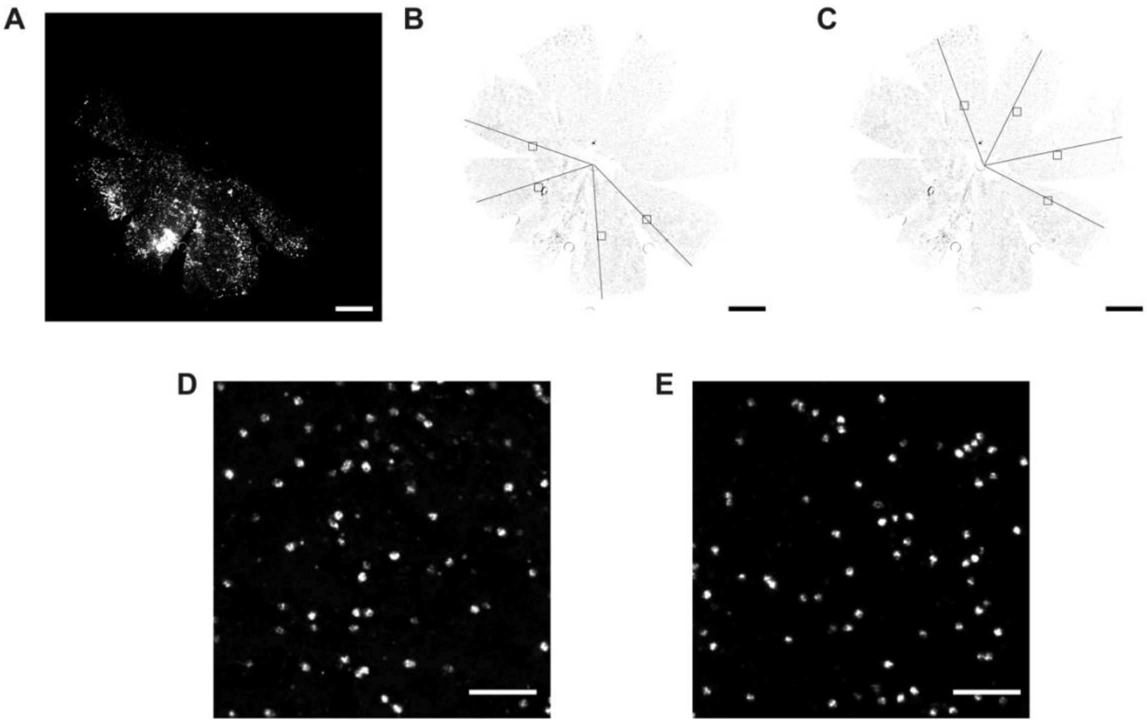


**Figure S1. Method for cone counting in rat retinal samples.** Representative retinae from a P180 S344ter rat subretinally injected with red fluorescent beads. Cones were stained using *in situ* hybridization for retinal cone arrestin-3 (*Arr3*). (A) Representative red fluorescent bead expression showing transduced (bead positive) and untransduced (bead negative) regions. (B) An ImageJ macro submitted *Arr3* stained retinal flatmount images to automatic processing and thresholding. Four lines were manually drawn from the periphery to the optic nerve head of the retina in the (B) transduced region and (C) untransduced region (A-C scale bar: 1 mm). 250 $\mu m^2$ boxes were automatically generated and placed along the midpoint of each line. *Arr3* stained cones in a single 250 $\mu m^2$ box in the (D) transduced and (E) untransduced region (scale bar: 50 $\mu m^2$). The mean number of cones in four 250 $\mu m^2$ boxes was calculated and used for statistical analyses.

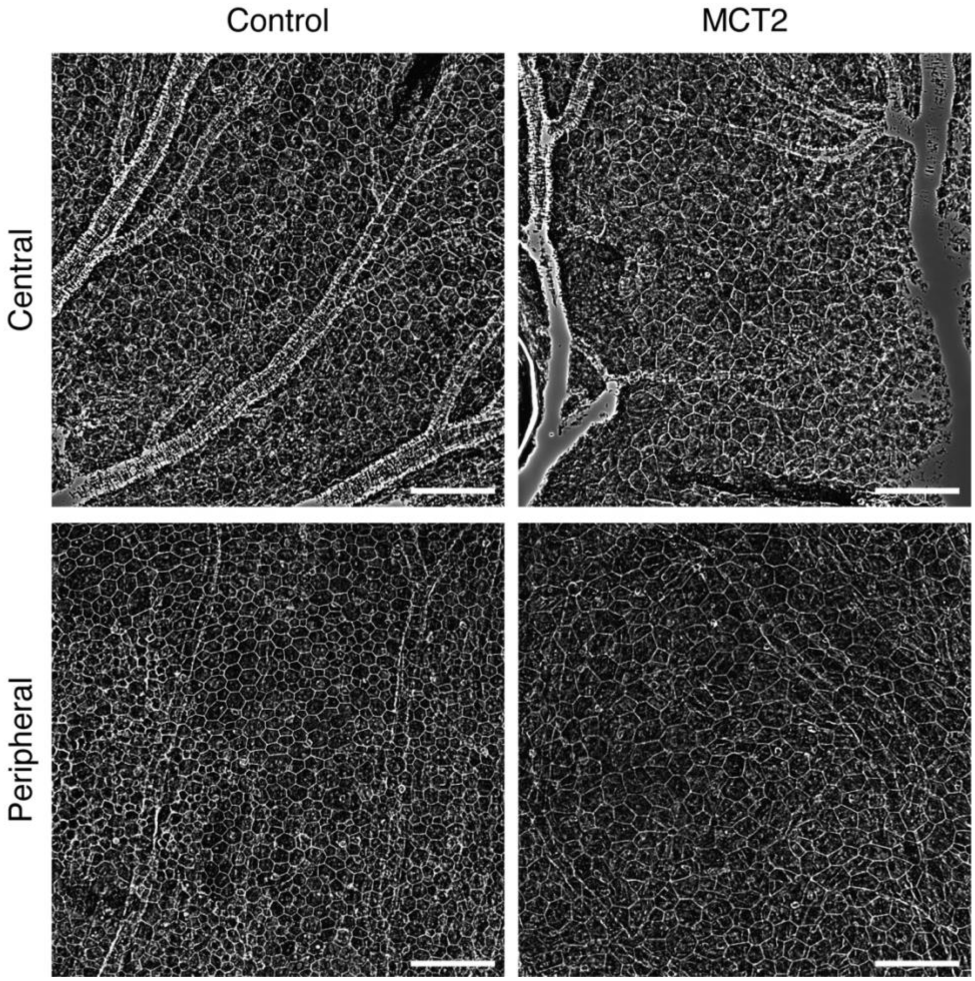


**Figure S2. Minimal disruption to wildtype rat RPE following subretinal injection of AAV8.Best1.MCT2.** Neonatal Sprague Dawley rats were subretinally injected with red fluorescent beads alone (control) or co-injected with AAV8.Best1.MCT2 (MCT2) (n=5). Representative images of P31 phalloidin stained RPE at the central and mid-peripheral region showing minimal changes to the RPE hexagonal structure and no other signs of toxicity (scale bar: 100 μm).

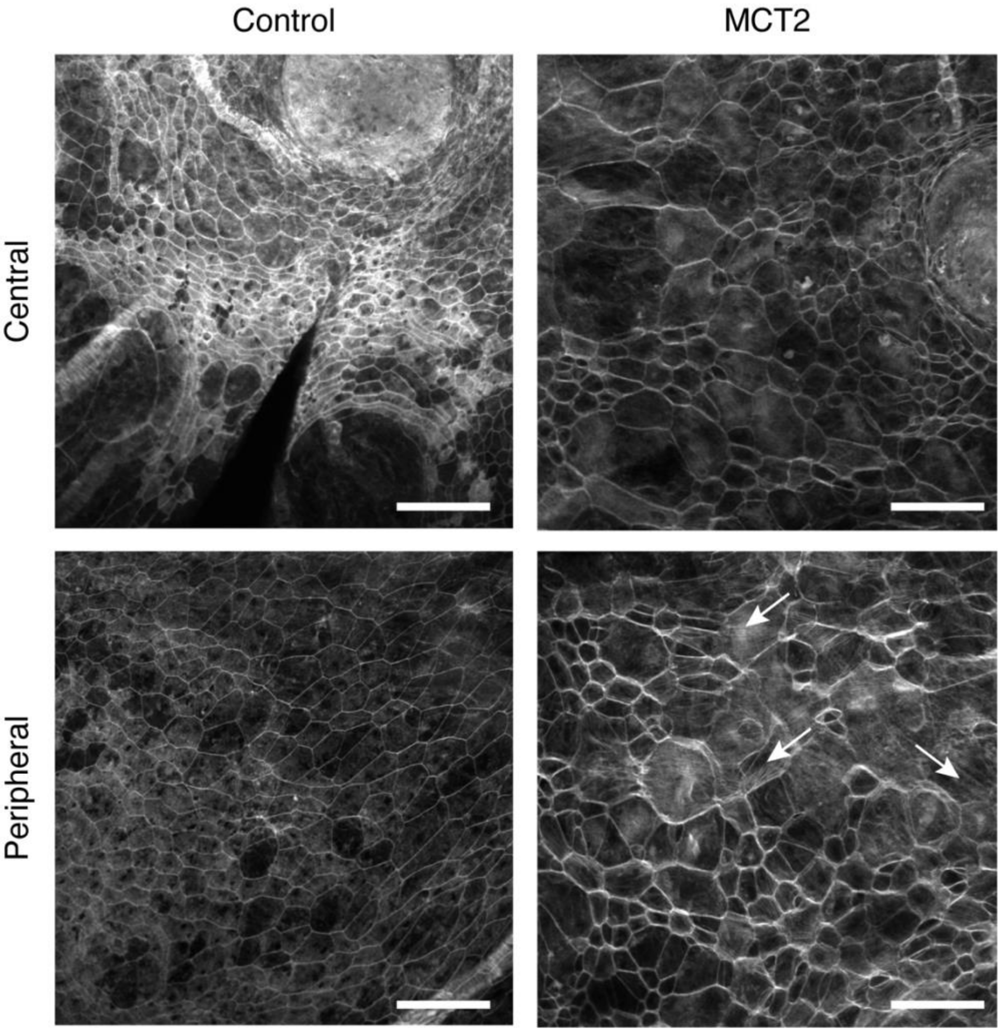


**Figure S3. Signs of toxicity in mouse FVB RPE following subretinal injection of AAV8.Best1.MCT2.** Neonatal FVB mice were subretinally injected with PBS (control) (n=5) or AAV8.Best1.MCT2 (MCT2) (n=7). Representative images of P40 phalloidin stained RPE at the central and mid-peripheral regions demonstrating increased disruption of RPE morphology and the upregulation of stress fibers (scale bar: 100 $\mu$m). Stress fibers are indicated with the white arrows.

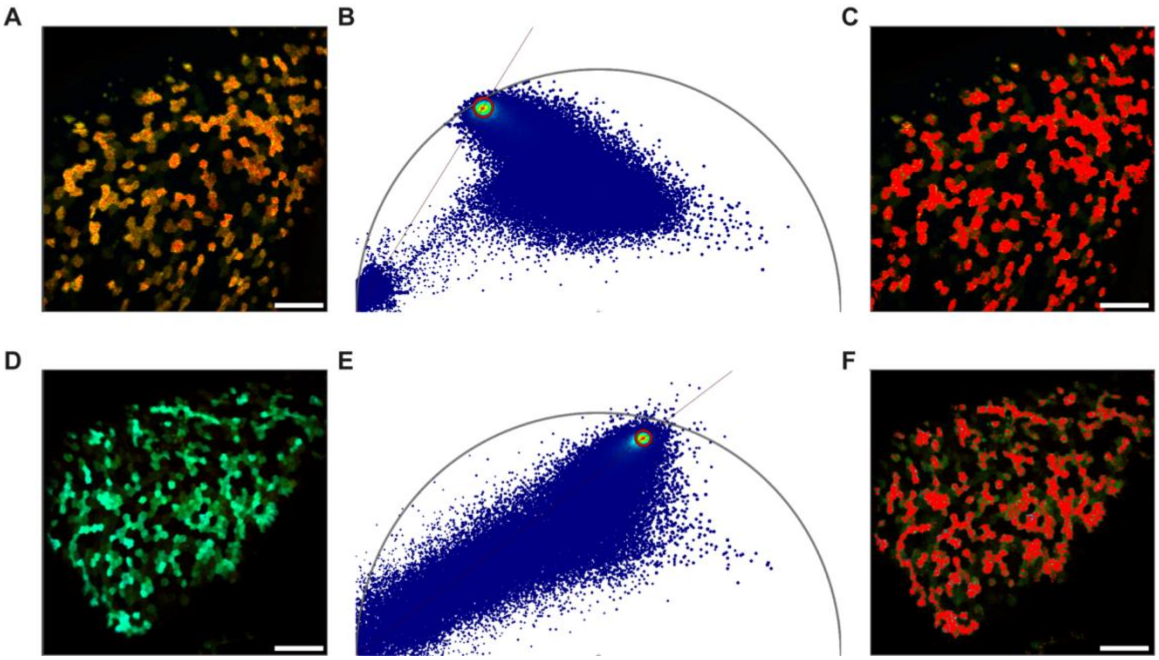


**Figure S4. FLIM phasor analysis.** RPE tissue from FVB mice subretinally injected with (A-C) AAV8.Best1.LiLac or (D-F) AAV8.Best1.GlucoSnFR-TS. (A&D) Representative lifetime image at baseline (scale bar: 100 $\mu$m). (B&E) The corresponding phasor plot demonstrating a 2D graphical view of lifetime distribution with each point on the plot corresponding to a pixel in the image. A single molecular species corresponding to (B) lactate or (E) glucose is selected within the red circle to calculate the lifetime of that sample. (C&F) Overlay in red demonstrating the subpopulation selected within the phasor plot.
